## Supplementary_figures_1_2 for "Dynamic layer-specific processing in the prefrontal cortex during working memory"

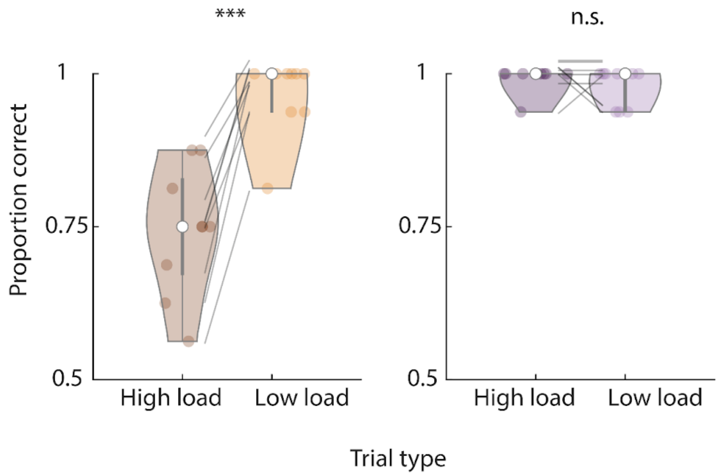


**Supplementary figure 1: Behavioral responses.** The left panel shows the accuracy of subjects in the high and low load trials. The right panel shows the accuracy of responding or abstaining to respond in motor trials.

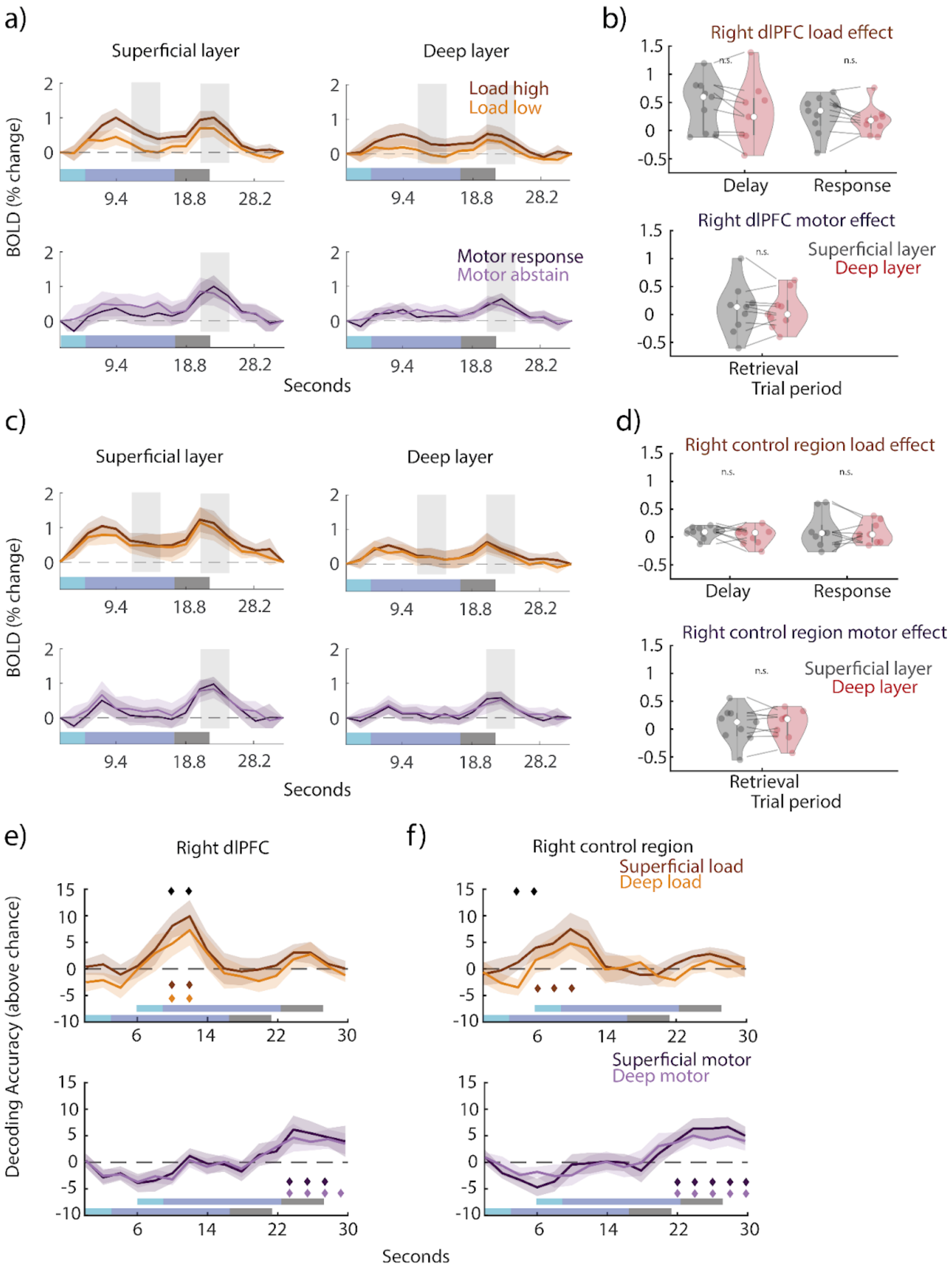

a)
Superficial layer
Deep layer
Load high
Load low
Motor response
Motor abstain
BOLD (% change)
Seconds
b)
Right dlPFC load effect
Delay
Response
Right dlPFC motor effect
Superficial layer
Deep layer
Retrieval
Trial period
c)
Superficial layer
Deep layer
BOLD (% change)
Seconds
d)
Right control region load effect
Delay
Response
Right control region motor effect
Superficial layer
Deep layer
Retrieval
Trial period
e)
Right dlPFC
f)
Right control region
Superficial load
Deep load
Superficial motor
Deep motor
Decoding Accuracy (above chance)
Seconds

**Supplementary figure 2: Layer-specific univariate BOLD time courses and multivariate decoding in the right hemisphere. a-d)** Same as **Fig. 2**, but for right dlPFC and right frontal control regions. Load effect between layers during delay in the right dlPFC: $p$ = 0.071, $t(8)$ = 2.08, $CI^{95}$ = [-0.0572, 0.3134], two-tailed t-test. Load effect between layers during the retrieval period in the right dlPFC: $p$ = 0.24, t = 1.27, $CI^{95}$ = [-0.123, 0.3042]. Motor effect between layers during retrieval in the right dlFPC: $p$ = 0.75, $t(8)$ = 0.332, $CI^{95}$ = [-0.1634,0.2037]. Load effect between layers during the delay period in the right control region: $p$ = 0.22, $t(8)$ = 1.339, $CI^{95}$ = [-0.0793, 0.2058]. Load effect between layers during the retrieval period in the right control region: $p$ = 0.66, t = 0.450, $CI^{95}$ = [-0.171,0.2309]. Motor effect between layers during retrieval in the right control region: $p$ = 0.99, $t(8)$ = 0.0104, $CI^{95}$ = [-0.121,0.1202]. **e-f)** Same as **Fig 3 a-b)**, but for right dlPFC and right frontal control regions. Right dlPFC superficial layer load decoding cluster at 10-12 s ($p$=0.0059, one-tailed permutation test); deep layer cluster at 10-12 s ($p$ = 0.045, one-tailed permutation test). Between layer load decoding cluster at 10-12 s ($p$ = 0.039, two-tailed permutation test). Right control region superficial layer load decoding cluster at 6-10 s ($p$ = 0.013, one-tailed permutation test). Between layer load decoding cluster at 4-6 s ($p$ = 0.030, two-tailed permutation test). Right dlPFC superficial layer motor decoding cluster at 24-28 s (p = 0.0013, one-tailed permutation test); deep layer cluster at 24-30 s ($p$ = 7.99e-04, one-tailed permutation test). Right control region superficial layer motor decoding cluster at 22-30 s ($p$ = 9.99e-05, one-tailed permutation test); deep layer cluster at 22-30 s ($p$ = 9.99e-05, one-tailed permutation test).
